# Identification of anthracnose (*Colletotrichum lentis*) race 1 resistance loci in lentil by integrating linkage mapping and a genome-wide association study

**DOI:** 10.1101/2021.03.16.435724

**Authors:** Tadesse S. Gela, Larissa Ramsay, Teketel A. Haile, Albert Vandenberg, Kirstin E. Bett

## Abstract

Anthracnose, caused by *Colletotrichum lentis*, is a devastating disease of lentil in Western Canada. Growing resistant lentil cultivars is the most cost-effective and environmentally friendly approach to prevent seed yield losses that can exceed 70%. To identify loci conferring resistance to anthracnose race 1 in lentil, biparental quantitative trait loci (QTL) mapping of two recombinant inbred line (RIL) populations was integrated with a genome-wide association study (GWAS) using 200 diverse lentil accessions from a lentil diversity panel (LDP). A major-effect QTL (*qAnt*1.*Lc*-3) conferring resistance to race 1 was mapped to lentil chromosome 3 and co-located on the lentil physical map for both RIL populations. Clusters of candidate nucleotide binding-leucine-rich repeats (NB-LRR) and other defense-related genes were uncovered within the QTL region. A GWAS detected 14 significant SNP markers associated with race 1 resistance on chromosomes 3, 4, 5, and 6. The most significant GWAS SNPs on chromosome 3 supported *qAnt*1.*Lc*-3 and delineated a region of 1.6 Mb containing candidate resistance genes. The identified SNP markers can be directly applied in marker-assisted selection to accelerate the introgression of race 1 resistance in lentil breeding.

## INTRODUCTION

Lentil (*Lens culinaris* Medik., 2n=2x=14) is an annual self-pollinating pulse crop with a genome size of ~4Gb (Arumuganathan and Earle, 1991). The crop is cultivated in more than 70 countries worldwide, with the production from Canada providing 40% of the world’s supply in recent years (2013-2019 average, FAOSTAT, 2021). Lentil provides an affordable source of dietary proteins, minerals, fiber, and carbohydrates and plays a vital role in alleviating malnutrition and micronutrient deficiencies (Srivastava and Vasishtha, 2012). Lentil production in the northern Great Plains of North America, particularly western Canada, is challenged by anthracnose caused by the fungus *Colletotrichum lentis* (Damm) (Damm et al., 2014). Yield loss of more than 70% can occur when susceptible cultivars experience high disease pressure (Chongo et al., 1999; Morrall and Pedersen, 1991). The disease has also been reported in Bangladesh, Bulgaria, Brazil, Ethiopia, Morocco, Pakistan, Syria, and USA and is considered of minor importance in other parts of the world (Bellar and Kebabeh, 1983; Bayaa and Erskine, 1997; Morrall, 1997; Kaiser et al., 1998).

Breeding and deployment of resistant cultivars is an important component of an integrated disease management strategy for preventing yield loss caused by anthracnose. Successful deployment requires continuous incorporation of new sources of resistance into elite breeding materials. Two pathogenic races of *C. lentis* (race 0 and race 1) have been described (Buchwaldt et al., 2004; Banniza et al., 2018). In cultivated lentil, no sources of high levels of resistance to the highly virulent race 0 have been found, however, resistance does occur in *Lens ervoides* in the tertiary gene pool (Tullu et al., 2006). Lentil accessions with resistance to anthracnose race 1 were identified in L. *culinaris* germplasm (Buchwaldt et al., 2004). Resistance to race 1 was transferred into elite breeding lines and resulted in the release of cultivars such as ‘CDC Robin’ and ‘CDC Redberry’ (Vandenberg et al., 2002, 2006). Since then, several lentil cultivars with partial resistance to race 1 have been released and deployed for production in Saskatchewan (Government of Saskatchewan, 2019).

Successful incorporation of race 1 resistance into improved lentil cultivars is possible through classical breeding but could be greatly improved if molecular breeding approaches could be employed. This requires use of molecular markers to identify the genes that control the quantitative traits; however, little is known about the causal genomic regions controlling race 1 resistance in lentil. Tullu et al. (2003) mapped race 1 resistance in lentil accession PI 320937 using RAPD markers and identified a major dominant gene and several minor genes. Segregation analysis of race 1 resistance in PI 320952 and PI 345629 revealed control by recessive and dominant genes (Buchwaldt et al., 2013).

More precise knowledge of accurate localization of QTL/genes and identification of linked molecular markers is an important step in development of effective MAS in lentil breeding. It also facilitates pyramiding of the resistance genes into lentil cultivars to achieve high levels of resistance against both races of anthracnose. Integration of QTL mapping in biparental populations and GWAS provides the technology to identify trait loci associated with resistance while refining the genomic regions with high resolution (Zhu et al., 2008). Both mapping strategies have been successfully used to identify QTL for multiple traits in lentil (biparental: Fedoruk et al., 2013, Ates et al., 2016; Aldemir et al., 2017; Subedi et al., 2018; Sari et al., 2018; Polanco et al., 2019; and GWAS: Khazaei et al., 2018; Kumar et al., 2018; Kumar et al., 2019).

In lentil, the low number and density of markers used for mapping studies have been the limiting factor for a long time (reviewed by Kumar et al., 2015). However, advances in next generation sequencing technologies have enabled the detection of large numbers of SNP sets from different marker genotyping platforms in lentil. Exome capture genotyping, which targets the genic regions of the genome, has been demonstrated to be an efficient method of high-throughput SNP discovery in lentil (Ogutcen et al., 2018). The SNP markers targeting the functional region of a genome may be of great importance to breeders because they are attributable to traits of interest under artificial selection through MAS. Here, we used exome capture genotyping on a diversity panel of lentil accessions and a lentil biparental RIL population (Haile et al., 2020). The objective of this research were: (i) to identify QTL for anthracnose race 1 resistance in two lentil RIL populations, (ii) to detect chromosomal regions associated with race 1 resistance using GWAS, and (iii) to compare the genomic regions detected in both mapping strategies to identify candidate genes involved in disease resistance.

## MATERIAL AND METHODS

### Plant materials

Two biparental-derived lentil RIL populations (LR-01 and LR-18) and a GWAS panel were used to identify loci conferring resistance to anthracnose race 1, LR-01 was derived from the cross ILL 1704 × CDC Robin (Haile et al., 2020); LR-18 was developed from the cross CDC Robin × 964a-46 (Tar’an et al., 2003). Both RIL populations were advanced to F7 by single seed descent before bulking and comprised a set of 102 and 139 RILs for LR-01 and LR-18, respectively. CDC Robin is a cultivar partially resistant to anthracnose race 1 and resistant to ascochyta blight (Vandenberg et al., 2002). Parents ILL 1704 and 964a-46 are susceptible to anthracnose race 1. The pedigree of breeding line 964a-46 includes ILL 5588, an ascochyta blight resistant landrace released as the cultivar Northfield in Australia (Ali, 1995). ILL 1704 is a landrace from Ethiopia with moderate resistance to ascochyta blight (Tullu et al., 2010). The GWAS panel was a subset of 200 lentil genotypes selected from the Lentil Diversity Panel (LDP; N=324) (Haile et al., 2020; Wright et al., 2021; http://knowpulse.usask.ca/Lentil-Diversity-Panel). The LDP consists of 324 accessions assembled from the gene banks of Plant Gene Resources of Canada (PGRC), the USDA, the International Center for Agricultural Research in the Dry Areas (ICARDA), and includes cultivars developed at the Crop Development Centre (CDC), University of Saskatchewan (USask).

### Disease phenotyping

*Colletotrichum lentis* isolate CT-21 representing race 1 (Banniza et al., 2018) was used to inoculate the RIL populations and GWAS panel. Fungal inoculum production and inoculation procedures were done as described by Gela et al. (2020). Disease reactions of the RIL populations and parents were evaluated in a growth chamber (GR48, Conviron, Winnipeg, Canada) environment at the USask College of Agriculture and Bioresources phytotron facility, Saskatoon, Canada. The GWAS panel was evaluated in a growth chamber and in an outdoor polyhouse. In the growth chambers evaluations, plants of each accession/RIL were grown in replicates, each consisting of sets of 38-cell cone trays (26.8 cm x 53.5 cm) per replication. The trays were filled with Sun Gro Horticulture Sunshine Mix LA4 (Sun Gro Horticulture, Bellevue, USA) and perlite (Specialty Vermiculite Canada, Winnipeg, MB) at 3:1 ratio (v/v). The susceptible control cultivar ‘Eston’ (Slinkard, 1981) and the RIL parental genotypes were included in each tray. Four weeks after seeding, plants were inoculated with a spore suspension (5 × 10^4^ spores mL^-1^) at 3 mL per plant using an airbrush. Plants were placed in an incubation chamber (relative humidity 90-100%) for 48h before being moved to misting benches (see Gela et al., 2020). The experiments were conducted separately for each population and arranged in randomized complete block design with five and seven replications for the RIL populations and the GWAS panel, respectively, which were blocked over time. For the RIL populations, two plants of each RIL were included in each of five sequential experimental runs and the mean disease severity score of the two plants per replicate was used for statistical analysis. For the GWAS panel, one plant of each accession was evaluated as an individual and repeated seven times. For GWAS accessions exhibiting segregation for disease reaction, an additional three runs were conducted to obtain representative disease scores. For all experiments the plants were scored for race 1 disease severity at 8-10 days post-inoculation (dpi) using a 0 to 10 rating scale with 10% increments.

The polyhouse experiment was conducted at the Department of Plant Sciences field laboratory at the USask. Four seeds of each accession and two seeds of Eston (susceptible control) were sown in individual 4.5 L pots (15.5 cm diameter) containing Sunshine Mix No. 4 (Sun Grow Horticulture^®^ Ltd., Vancouver, BC, Canada). The plants were grown under open field conditions for 6 weeks (early flowering stage), then immediately before inoculation, polyhouse tunnel covers of translucent thin plastic sheeting were installed by suspending them 1.5 m above the ground over the pots. The area under the cover was equipped with a misting irrigation system. Each pot was sprayed with approximately 36 mL (6 mL plant ^-1^) of aqueous spore suspension (5 × 10^4^ spores mL^-1^) of race 1 isolate CT-21 using a pressurized knapsack sprayer. The inoculations were performed in the evening to avoid high temperature conditions and to facilitate the germination of spores on the leaves. To promote disease development after inoculation, misting irrigation was applied from early morning to evening for 30 s every 15 min. The experiment was conducted in a randomized complete block design (RCBD) with three replications. Disease severity data were collected 14 d after inoculation using a 0 to 10 rating scale with 10% increments. Data were converted to percentage disease severity using the class midpoints for data analysis.

### Statistical analysis of phenotypic data

Analysis of variance (ANOVA) for the phenotypic data were performed using the PROC GLIMMIX procedure of SAS v.9.4 (SAS Institute, Cary, USA). Broad-sense heritability (h_B_^2^) of single and combined environments disease severity scores of the GWAS panel were calculated with the lme4 package (Bates et al., 2015) in R (R Core Team, 2020) using the equation: σ^2^_G_/(σ^2^_G_ + σ^2^e/r) and σ^2^_G_/[(σ^2^_G_) + (σ^2^_GE_/_E_) + (σ^2^e/Er)], where σ^2^_G_ is the genotypic variance, σ^2^_GE_ is variance of the genotype × environment interactions, σ^2^e is the error variance, r is the number of replications in each environment and E is the number of environments (Knapp et al., 1985). Lsmeans of disease severity scores were calculated for each environment and for combined environments and subjected to square root transformation to improve the normality of the skewed distribution to perform GWAS analysis as described by Li et al. (2019). Spearman’s rank correlation of disease severity between test environments was performed using the procedure CORR in SAS. For RIL populations, mean disease severity data were calculated from the replicates and used for QTL mapping.

### SNP genotyping

Genomic DNA was isolated from fresh leaves of 2 to 3-week-old seedlings of the GWAS panel and LR-01 populations, including the parents (Haile et al., 2020). All lines were genotyped with a custom exome capture assay using protocols previously described by Ogutcen et al. (2018). The SNPs were called using lentil reference genome (CDC Redberry genome assemble *v2.0;* Ramsay et al., 2019). The resulting variant call format (VCF) file was filtered using VCFtools (Danecek et al., 2011) to retain SNPs with less than 10% missing allele calls (minimum read depth=5) and SNPs with minor allele frequency greater than 5%.

### Linkage map construction and QTL mapping

*LR-01 population*. A draft genetic linkage map consisting of 21,634 SNPs grouped into seven linkage groups corresponding to the seven haploid lentil chromosomes was generated using the MSTMap software (Wu et al., 2008). This high-density genetic map was further subjected to bin grouping of the redundant SNP markers using BIN functionality employed in QTL ICIMapping 4.1 software (Meng et al., 2015). A marker representing each bin was retained on the map, and the map distances in centimorgans (cM) between markers were calculated using the Kosambi function (Kosambi, 1944). All genotyping information for the LR-01 genetic linkage map can be found through the KnowPulse database accessible at: https://knowpulse.usask.ca/Geneticmap/2695342 (accessed January 6, 2021).

*LR-18 population*. The genetic linkage map developed earlier by Fedoruk et al. (2013) using the SNPs assayed using a 1536-SNP Illumina Golden Gate array (Illumina, San Diego, CA) was used for QTL mapping. The map consisted of 550 SNP markers, seven SSR markers, and four morphological markers. The total map distance was 697 cM with an average marker distance of 1.2 cM. All genotyping information for the LR-18 genetic linkage map can be found through the KnowPulse database accessible at: https://knowpulse.usask.ca/Geneticmap/2686922 (accessed January 13, 2021).

The QTL analyses were performed using R/qtl software (http://www.rqtl.org/; Broman et al., 2003). The QTL genotype probabilities were calculated along the chromosome at 1 cM intervals assuming a genotyping error rate of 1.0e^-4^ and using the Kosambi map function (Kosambi 1944). Multiple QTL mapping was completed with the *stepwiseqtl* function (Broman et al., 2003) using Haley-Knott regression (Haley and Knott, 1992). The optimal QTL model was chosen based on the highest penalized LOD score (Manichaikul et al., 2009) after forward and backward selection and elimination modelling using *stepwiseqtl* function. Penalties for model selection and genomewide significance threshold (α = 0.05) were determined by 1000 permutations with *scantwo* function for two-dimensional QTL scan. The confidence intervals for each QTL were estimated using the “*lodint*” function that calculated the 1.5 LOD support intervals. The percentage of the phenotypic variance explained (PVE) and effects of QTLs were obtained by fitting a mixed linear model using the “*fitqtl*” function (Broman et al., 2003).

### Genome-wide association analysis

The population structure of the association mapping panel was assessed using a pruned subset of 6,516 unlinked SNP markers generated after removing SNPs with minor allele frequency of <10% and linkage disequilibrium (r^2^ <0.2) at a sliding window of 1 Mb using SNPRelate package (Zheng et al., 2012). The Bayesian model-based clustering implemented in STRUCTURE V2.3.4 (Pritchard et al., 2000) was used to estimate the number of subpopulations (K). The number of K sets from K=2 to K=10, with 10 times independent runs for each K, 50,000 burn-in iterations, and 100,000 Markov chain Monte Carlo sampling replicates were conducted. The optimal K-values were determined using STRUCTURE HARVESTER (Earl & vonHoldt, 2012), and visualized by STRUCTURE PLOT (Ramasamy et al., 2014). Genotypes with membership probabilities <60% were considered admixtures (Falush et al., 2003). Principal component analysis (PCA) (Patterson et al., 2006) and the genetic kinship matrix were conducted using the Genomic Association and Prediction Integrated Tool (GAPIT) (Lipka et al., 2012).

Marker-trait associations (MTA) were tested using a compressed mixed linear model (CMLM) (Zhang et al., 2010) including the population structure (Q) and kinship (K). Association tests were run using the software GAPIT implemented in R software (Lipka et al., 2012). The quantile-quantile (Q-Q) plot’s fattiness was inspected to compare the results from Q + K (population structure and kinship), and PC + K (principal component and kinship) analysis. However, because the Q-Q plots generated using the two approaches were similar, only the results of the PC+K analysis are presented (Supplemental Figure S3). SNPs with -Log_10_(p-value) ≥ 5.2 were considered to have significant associations based on a Bonferroni threshold (1/n) correction at p > 6.6×10^-6^. Manhattan and Q-Q plots were generated with the R package qqman (Turner, 2014).

### Candidate gene analysis

The physical location of markers that defined the QTL intervals were identified using the CDC Redberry genome assembly v.2.0 (Ramsay et al., 2019) and examined for the presence of candidate genes associated with disease resistance. The annotated genes identified were used in BLAST analysis of GenBank (NCBI) database to confirm their functions in other plant species. All reported disease resistance (R-) or defense-related genes in plants were considered for identification of candidate genes.

## RESULTS

### Phenotypic variation

In both RIL populations, the resistant parent CDC Robin had a resistant reaction with mean disease severity of 23%. The susceptible parents, ILL 1704 and 964a-46 had susceptible reactions with the mean disease scores of 95% and 93%, respectively. Most of the RILs exhibited disease severity between the range of their parental lines, but some even exhibited lower disease severity than CDC Robin (Figure 1 a and b). The disease reactions of the two RIL populations to anthracnose race 1 ranged from 5 to 95%, with an overall average mean of 53.6% for LR-01 and 54.2% for LR-18. The RILs in both populations had bimodal frequency distributions which fitted a 1:1 resistant to susceptible segregation ratio, indicating monogenic segregation for resistance to anthracnose race 1 (Supplemental Table S1).

**Figure 1.**
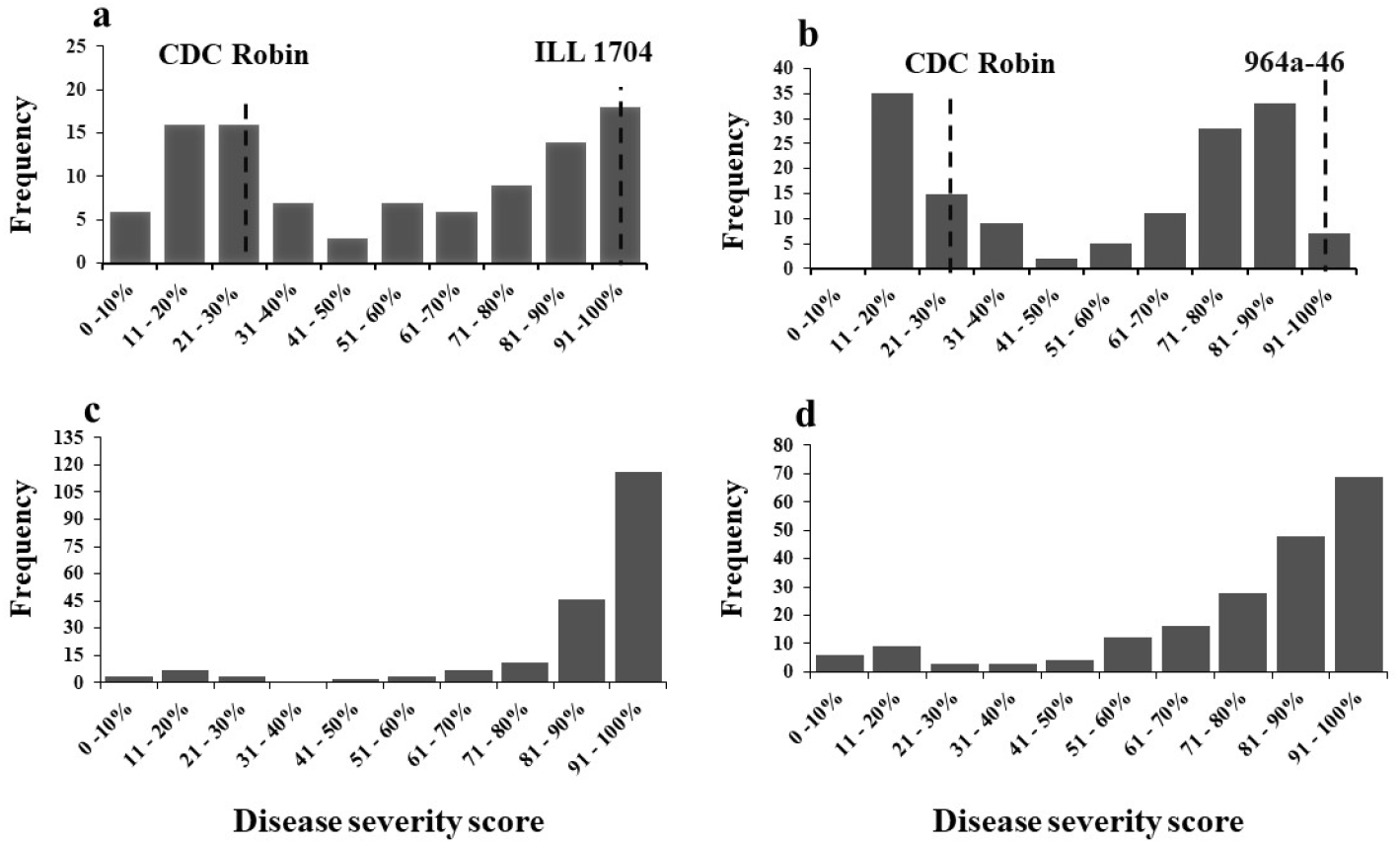
Frequency distribution of anthracnose race 1 severity of; (a) 102 RILs of LR-01 (ILL 1740 × CDC Robin), and (b) 139 RILs of LR-18 (CDC Robin × 964a-46) in growth chamber conditions. (**c, d**) frequency distributions of the 200 lentil genotypes in the GWAS panel evaluated under growth chamber and polyhouse conditions. The vertical lines indicate the average disease severity of the parents. Disease severity was rated on a 0-10 scale, where the disease severity score increased in 10% increments.

The disease severity distribution for the GWAS panel was skewed towards susceptibility in both testing environments (Figure 1 c and d). Under growth chamber conditions, 6.5, 8.5, and 85%; and for polyhouse conditions, 9, 25.5 and 65.5% of the genotypes had resistant, intermediate, and susceptible reactions, respectively. The results suggest the presence of limited sources of resistance among most of the genotypes tested against race 1, even though it was less aggressive than *C. lentis* race 0 (Buchwaldt et al., 2004). The differences in disease severity scores among the genotypes in each environments and genotype × environment interactions were significant (Table 1). The estimated broad-sense heritability was high, 0.96 for growth chamber, 0.88 for polyhouse and 0.92 for combined analysis of both environments (Table 1), demonstrating that race 1 resistance was controlled by genetic factors and that the data could be used for accurate mapping of race 1 resistance genes. A positive correlation (Spearman r = 0.61, p < 0 .001) was observed between growth chamber and polyhouse data for race 1 response of genotypes.

**Table 1.** Analysis of variance components for anthracnose race 1 severity of 200 lentil genotypes evaluated under growth chamber and polyhouse conditions.

| Environment | Mean | Range | $h_B^2$ | $\sigma^2_G$ | $\sigma^2_{GE}$ | $\sigma^2_e$ |
| --- | --- | --- | --- | --- | --- | --- |
| Growth chamber | 84.2 | 6.4 - 95 | 0.96 | 405.1*** | - | 104.8 |
| Polyhouse | 76.0 | 5.0 - 95 | 0.88 | 454.5*** | - | 180.5 |
| Combined | 80.1 | 5.7 - 95 | 0.92 | 382.8*** | 45.55*** | 133.2 |

\*\*\* $p < 0.0001$ , $\sigma^2_G$ , genotypic variance; $\sigma^2_{GE}$ , genotype × environment variance; $\sigma^2_e$ , error variance; $h_B^2$ , heritability.

### Linkage map construction for the LR-01 population

Seven linkage groups (LG) were constructed from the segregation patterns of 21,634 SNPs. LGs were assigned to their respective chromosomes based on where markers lie in the reference genome. The detailed information for the LR-01 linkage map is provided in supplemental Table S2. The number of SNPs assigned to each linkage group (LG) ranged from 1710 SNP markers (LG 7) to 5120 (LG 2). After binning of SNPs with redundant information, the 21,634 SNPs were grouped into 921 recombination bins and 1807 single markers. In total, 2728 informative loci distributed along the seven LGs were retained on the LR-01 linkage map. The map covered a total length of 1643.8 cM, with an average distance between the neighboring SNP markers of 0.6 cM. The length of each LG varied from 164.6 cM for LG 7 to 299.5 cM for LG 3.

### QTL mapping of anthracnose race 1 resistance

A single significant QTL conferring resistance to anthracnose race 1 was identified on chromosome 3 using both LR-01 and LR-18 RIL populations and designated as *qAnt*1.*Lc*-3 (Figure 2). On the LR-18 linkage map, the QTL was flanked by SNP markers LcC03673p249/LcC03441p105 and LcC09426p518, with an interval ranging from 43.2 to 51.5 cM. The SNP markers were mapped within a 7.6 Mb (30704841 to 38275723 bp) physical interval on the CDC Redberry genome assembly v.2.0 (Ramsay et al., 2019). For the LR-01 population genetic map, *qAnt*1.*Lc*-3 mapped to an interval of 53.2 cM to 70.8 cM corresponding to a physical location of 29313444 to 38758654 bp (9.4 Mb region) on chromosome 3. Importantly, the high-density genetic map of the LR-01 population contains a number of SNP markers associated with race 1 resistance in the interval of the QTL region. Among these, a cluster of markers mapped to Lcu.2RBY.unitig0289 which is unanchored in the reference genome, suggesting it should be part of chromosome 3. The percentage of variation in race 1 resistance explained by *qAnt*1.*Lc*-3 varied from 66.6 to 69.8%, with a LOD score value ranging from 24.3 to 37.1 (Table 2). As expected, CDC Robin (the resistant parent) contributed the resistance allele for *qAnt*1.*Lc*-3.

**Figure 2.**
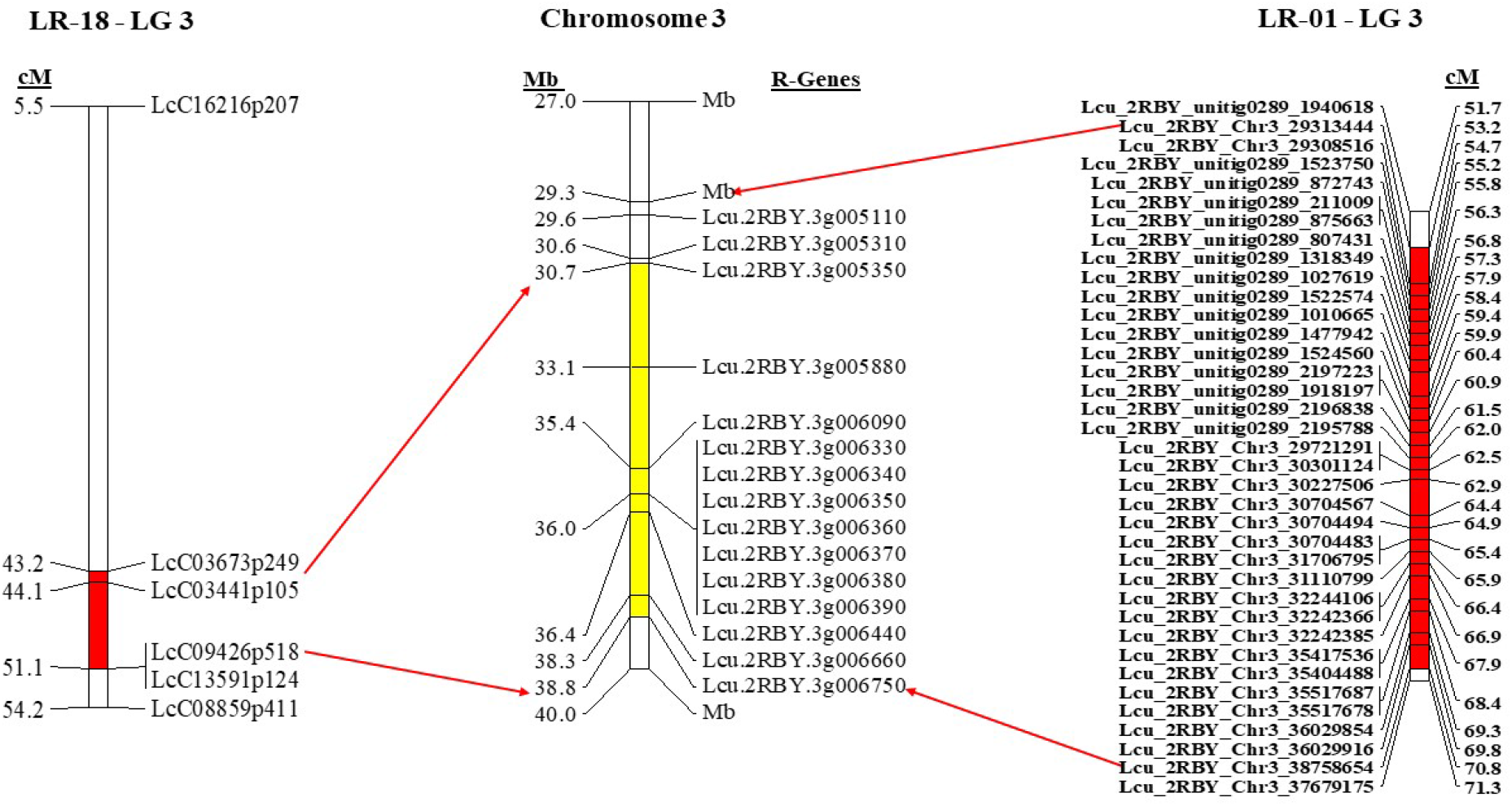
Position of anthracnose race 1 resistance QTL on linkage group (LG) 3 of the recombinant inbred line (RIL) populations LR-18 (CDC Robin × 964a-46; left) and LR-01 (ILL 1704 × CDC Robin; right) evaluated under growth chamber conditions. The red regions on the bar highlights the QTL interval on LGs; and the yellow region depicts the interval overlapped for both LGs on lentil chromosome 3 (*Ref. genome* v2.0) and the predicted candidate disease resistance genes (R-genes) is on the right. The positions are in centimorgan (cM) and mega base pairs (Mb) as indicated on the top of the bar

**Table 2.**
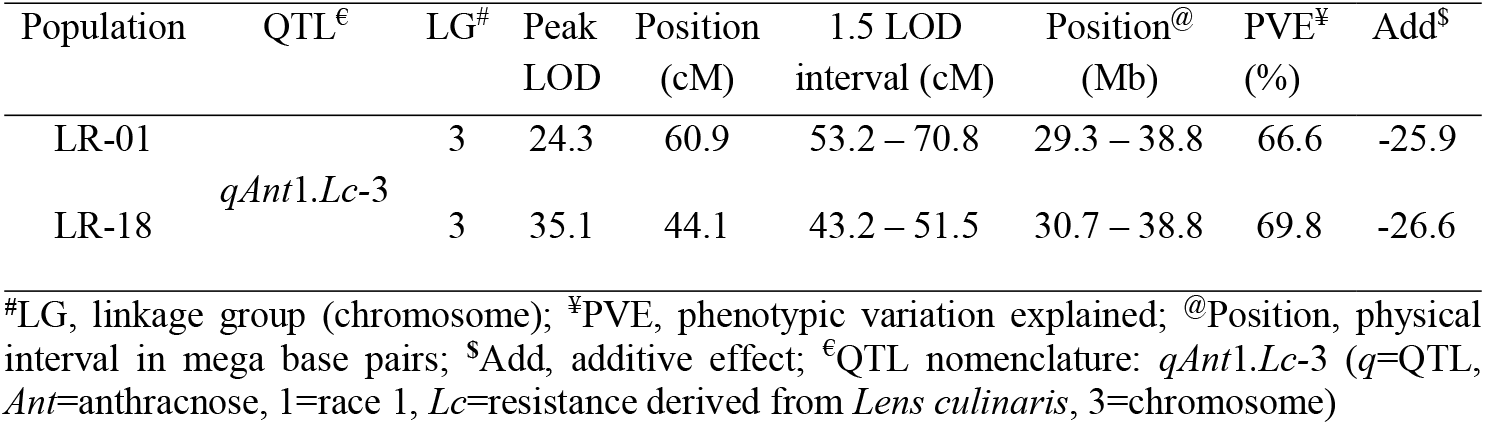
QTL mapping of anthracnose race 1 resistance detected by multiple QTL models of R/qtl in two biparental RIL populations: LR-01 (ILL 1704 × CDC Robin) and LR-18 (CDC Robin × 964a-46).

### Genome-wide association study of anthracnose race 1 resistance

A total of 152,011 SNP markers were used for marker-trait association analysis. The number of SNP markers per chromosome varied from 16,848 on chromosome 7 to 26,349 on chromosome 2 (Supplemental Figure S1). The average physical distance between two markers used in this study was approximately 26 kb, across a genome size of ~4 Gb with a mean of 21,716 SNP markers per chromosome. The SNP markers were evenly distributed and adequately covered the genome for the purpose of GWAS analysis.

The model-based population structure analysis revealed that the 200 lentil genotypes could be grouped into three major subpopulations, and that finer hierarchical structures were evident in the diversity panel (Supplemental Figure S2 a). Using k = 3, 90.5% of the accessions were assigned to three groups, and only 9.5% of the accessions were assigned to mixed populations (Supplemental Figure S2 b). The principal component analysis (PCA) showed that the variance explained by the eigenvalue of each principal component (PC) dropped rapidly after the first three PCs, which explained approximately 35% of the total genetic variances for the diversity panel (Supplemental Figure S2 c). The results of the PCA were consistent following STRUCTURE analyses (Supplemental Figure S2 d). Thus, the first three PCAs were used as a covariate in the mixed linear model in the GWAS analysis and were also confirmed by visual inspection of Q-Q plots (Supplemental Figure S3).

GWAS analysis using the combined disease severity data detected 14 SNPs that were significantly associated with race 1 resistance in the lentil genome (-Log_10_^(p)^ ≥ 5.2) (Figure 3; Table 3). The GWAS analysis for individual environments identified 26 and 11 SNPs associated with race 1 resistance under growth chamber and polyhouse conditions, respectively (Figure 3 and Supplemental Table S3). Most of the loci detected were common between both environments with varying p-values of SNPs associated with the loci. The phenotypic variation (R^2^) explained by an individual significant SNP marker ranged from 58 to 69%.

**Figure 3.**
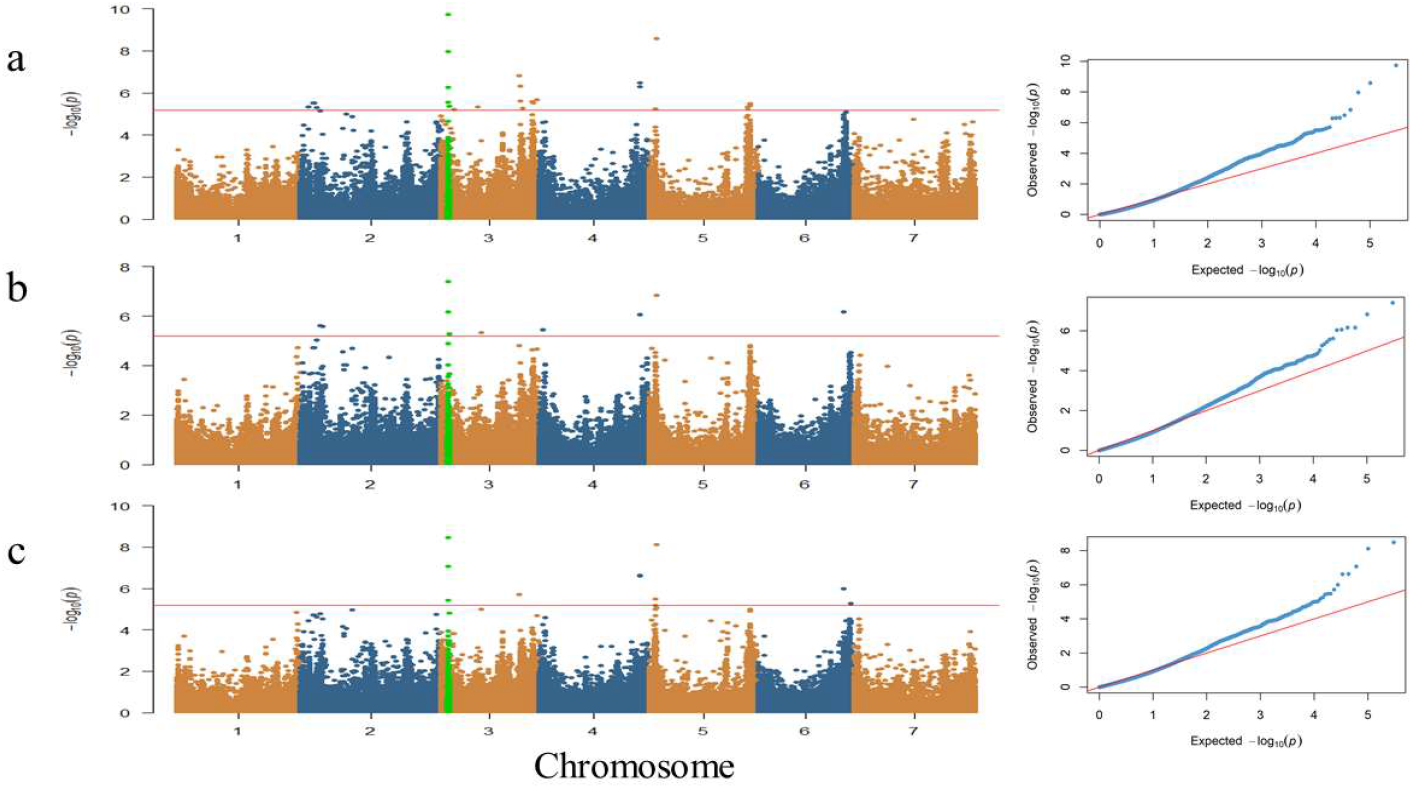
Manhattan and Quantile-quantile (Q-Q) plots of genome wide association study (GWAS) for anthracnose race 1 resistance in 200 lentil accessions evaluated in (**a**) the growth chamber, (**b**) the polyhouse and (**c**) the combined lsmean of disease severity scores from both environments. Alternating colors indicate the 7 chromosomes and the Y-axis indicates -log_10_ of p-values with significant association at 5.2 (red line). The green dots on chromosome 3 represent the SNP marker in the QTL (qAnt1.Lc-3) interval from biparental populations.

The SNPs previously associated with the *qAnt*1.*Lc*-3 region on chromosome 3 were again the most significant with -log_10_ (*p*) = 9.6 and R^2^ = 69% (Figure 3 and Table 3) and were detected in all analyses. This region was tagged with four significant SNP markers (Lcu.2RBY.Chr3.33827173, Lcu.2RBY.Chr3.33827185, Lcu.2RBY.Chr3.34117023 and Lcu.2RBY.Chr3.35384298) and spanned 1.6 Mb. Two other genomic regions on chromosome 3 displayed a peak SNP marker at 341.3 Mb (R^2^ = 66%) and 417.9 Mb (R^2^ = 65%) for combined and growth chamber analysis. Two SNP markers associated with race 1 resistance were detected on chromosome 4 within 4 bp (442702129 bp to 442702133 bp) in all analyses. Two other significant regions were identified for combined and growth chamber data on chromosome 5 from 28.4 to 33.7 Mb for all analysis and from 427.5 to 437.9 Mb. One SNP marker located at 374.3 Mb on chromosome 6 was detected for the combined and the polyhouse analyses (Table 3).

**Table 3.** SNP markers associated with anthracnose race 1 resistance using combined lsmean data of disease severity from growth chamber and polyhouse for a set of 200 lentil accessions.

| SNP Marker | Chr | Position (Mb)<sup>#</sup> | P.value | MAF | R<sup>2</sup><sup>\$</sup> |
| --- | --- | --- | --- | --- | --- |
| Lcu.2RBY.Chr3.33827173 | 3 | 33827173 | 1.38E-06 | 0.16 | 0.65 |
| Lcu.2RBY.Chr3.33827185 | 3 | 33827185 | 2.43E-08 | 0.15 | 0.67 |
| Lcu.2RBY.Chr3.34117023 | 3 | 34117023 | 2.47E-10 | 0.14 | 0.69 |
| Lcu.2RBY.Chr3.35384298 | 3 | 35384298 | 3.24E-06 | 0.14 | 0.65 |
| Lcu.2RBY.Chr3.341261994 | 3 | 341261994 | 4.30E-07 | 0.23 | 0.66 |
| Lcu.2RBY.Chr3.417940994 | 3 | 417940994 | 6.13E-06 | 0.06 | 0.64 |
| Lcu.2RBY.Chr4.442702129 | 4 | 442702129 | 8.32E-08 | 0.11 | 0.66 |
| Lcu.2RBY.Chr4.442702133 | 4 | 442702133 | 8.48E-08 | 0.11 | 0.66 |
| Lcu.2RBY.Chr5.28582530 | 5 | 28582530 | 3.74E-06 | 0.12 | 0.65 |
| Lcu.2RBY.Chr5.28637458 | 5 | 28637458 | 3.74E-06 | 0.12 | 0.65 |
| Lcu.2RBY.Chr5.33721990 | 5 | 33721990 | 1.82E-09 | 0.21 | 0.68 |
| Lcu.2RBY.Chr5.437910070 | 5 | 437910070 | 3.23E-06 | 0.13 | 0.65 |
| Lcu.2RBY.Chr5.437944230 | 5 | 437944230 | 3.95E-06 | 0.13 | 0.65 |
| Lcu.2RBY.Chr6.374326758 | 6 | 374326758 | 8.88E-07 | 0.11 | 0.65 |

<sup>#</sup>Physical positions, <sup>\$</sup>Explained phenotypic variance per marker

### Candidate gene prediction

We explored the candidate genes associated with anthracnose race 1 resistance in the region of *qAnt*1.*Lc*-3 physical genomic intervals that overlapped for both genetic maps and the GWAS. The interval covered 8 Mb from 30.0 - 38.0 Mb on chromosome 3 and contains 119 annotated genes. Among these, 11 genes encode for typical resistance (R) genes, which include nucleotide-binding-leucine-rich repeat (NB-LRR) domain disease resistance proteins (Table 4), and 46 are known to be involved in defense response reactions to pathogens and other stresses (Supplemental Table S4). Moreover, two NB-LRR domain genes (Lcu.2RBY.L001220 and Lcu.2RBY.L001240) were tagged by SNPs in unitig Lcu.2RBY.unitig0289 but mapped in the QTL region of the LR-01 genetic map, providing further evidence that they possibly correspond to a chromosome 3 NB-LRR domain cluster. Within the *qAnt*1.*Lc*-3 region, the most significant GWAS SNPs include one located 103 kb upstream of the gene encoding an anthranilate N-benzoyltransferase protein (Lcu.2RBY.3g005860), two significant SNPs within genes involved with cellulose synthase (Lcu.2RBY.3g005880), and one significant SNP located within a gene encoding for a disease resistance protein TIR-NBS-LRR domain (Lcu.2RBY.3g006090). Almost all the significant GWAS SNPs identified are located within or close to an annotated gene (Supplemental Table S4).

**Table 4.** A subset of candidate resistance genes associated with anthracnose race 1 resistance identified in the interval of QTL and GWAS regions according to gene annotation.

| **Chr<sup>s</sup>** | **Gene ID** | **Physical position (bp)<sup>#</sup>** |  | **Annotation** |
| --- | --- | --- | --- | --- |
| Chr3 | Lcu.2RBY.3g005310 | 30641401 | 30646173 | NB-ARC domain disease resistance protein |
| Chr3 | Lcu.2RBY.3g005350 | 30735946 | 30736590 | Wall-associated receptor kinase-like protein |
| Chr3 | Lcu.2RBY.3g005880<sup>ß</sup> | 33819256 | 33828131 | Cellulose synthase |
| Chr3 | Lcu.2RBY.3g005910<sup>ß</sup> | 34117126 | 34118497 | Anthranilate N-benzoyltransferase |
| Chr3 | Lcu.2RBY.3g006090<sup>ß</sup> | 35383081 | 35387584 | TIR-NBS-LRR class disease resistance protein |
| Chr3 | Lcu.2RBY.3g006330 | 35972988 | 35973704 | LRR & NB-ARC domain disease resistance protein |
| Chr3 | Lcu.2RBY.3g006340 | 35974000 | 35976197 | LRR & NB-ARC domain disease resistance protein |
| Chr3 | Lcu.2RBY.3g006350 | 35976249 | 35981698 | LRR & NB-ARC domain disease resistance protein |
| Chr3 | Lcu.2RBY.3g006360 | 35981709 | 35982218 | NB-ARC domain disease resistance protein |
| Chr3 | Lcu.2RBY.3g006370 | 35982494 | 35982865 | CC-NBS-LRR resistance protein |
| Chr3 | Lcu.2RBY.3g006380 | 36028314 | 36032907 | LRR & NB-ARC domain disease resistance protein |
| Chr3 | Lcu.2RBY.3g006390 | 36037105 | 36044777 | LRR & NB-ARC domain disease resistance protein |
| Unitig0289 | Lcu.2RBY.L001220 | 1917198 | 1921007 | LRR & NB-ARC domain disease resistance protein |
| Unitig0289 | Lcu.2RBY.L001240 | 1939874 | 1942161 | NB-ARC domain disease resistance protein |

<sup>#</sup> Position in bp according to CDC Redberry genome ref. v.2.0, <sup>ß</sup>Genes identified with GWAS SNPs, <sup>s</sup>Chromosome

## DISCUSSION

Developing host resistance is the most preferable, economical, and sustainable strategy for managing anthracnose in lentil production. However, the process requires sufficient information about genetic sources of resistance and identification of resistance loci associated with races of anthracnose to use marker assisted breeding strategy for gene pyramiding. We used a combination of traditional QTL mapping and GWAS to increase our understanding of anthracnose race 1 resistance in lentil. To our knowledge, this study is the first on anthracnose race 1 resistance in lentil to employ biparental QTL mapping and GWAS using a large number of physical SNP marker positions.

Analysis of the differential responses of the RIL populations and GWAS accessions to race 1 inoculation provided frequency distributions and heritability estimates of the inheritance of disease resistance. The disease reactions of the RILs of both populations displayed a bimodal distribution that fit a 1:1 ratio indicating a Mendelian one gene model, indicative of a major QTL identified in the populations. Analysis of variance for the GWAS panel revealed significant variation among genotypes. However, disease reaction frequencies of the lines showed a skewed distribution to greater susceptibility. The narrow genetic base for anthracnose resistance is evident in current collections of lentil accessions from different gene banks. For example, previous evaluations of resistance to anthracnose race 1 of lentil accessions from 50 countries identified 16 (0.9%) of 1771 (Buchwaldt et al., 2004), and 15 (2.6%) of 579 (Shaik et al., 2013). This may be attributable to genetic bottlenecks that were created at the time of domestication (Sonnante et al., 2009) due to the absence of the disease in its region of domestication. Until recently, the disease was considered minor or had not been reported in other parts of the world (Banniza et al., 2018), including its center of origin and/or diversity where lentil has been grown for centuries. Conversely, the rapid expansion of the lentil crop in the prairie ecosystem, particularly in the Western Canadian production system, coincides with the incidence of anthracnose (Morrell, 1997). Tanksley and McCouch (1997) argued expansion of a few improved cultivars into a modern agricultural system can result in emergence of new disease threats that can occur due to plant pathogens host shift (Silva et al., 2012).

Breeding for partial resistance to anthracnose race 1 in lentil started with release of the cultivar CDC Robin (Vandenberg et al., 2002), which was derived from PI 320952 (Indianhead). The resistance in PI 320952 was shown to be governed by a recessive and a closely linked dominant gene (Buchwaldt et al., 2013). In this study, we mapped a major QTL (qAnt1.Lc-3) associated with race 1 resistance from CDC Robin on chromosome 3 in both populations. The QTL *qAnt*1.*Lc*-3 accounts for 66.6 −69.8% of the variance in resistance to race 1. Bhadauria et al. (2017) reported race 1 resistance QTL on chromosomes 2, 3, and 5 of the *L. ervoides* genome. We do not have sufficient information to determine if the QTL on chromosome 3 of *L. ervoides* is the same as that in the cultivated lentil but both genomes are largely collinear along their length of this chromosome (Ramsay et al., *unpublished data*). Large portions of the physical interval of *qAnt*1.*Lc*-3 were colocalized for the LR-01 and LR-18 population genetic maps, indicating a strong association of the genomic region with race 1 resistance.

The association mapping approach is suited for the detection of high-resolution QTLs, as it captures a larger portion of the recombination events that have accumulated inside an association panel (Zhu et al., 2008). Exploration of high throughput marker data makes GWAS more efficient and provides a rapid method to identify significant genomic regions associated with traits of interest for candidate gene prediction (Yano et al., 2016). In the current study, marker-trait associations identified 14 SNPs associated with race 1 resistance on chromosomes 3, 4, 5 and 6. Most of these MTA were consistent across test conditions, explained by the high heritability estimates and the high correlation observed between results from growth chamber and polyhouse environments. Chongo and Bernier (1999) also reported significant correlation between field inoculations and anthracnose screening under controlled conditions for resistant *L. culinaris* germplasm. The major locus on chromosome 3 identified using the biparental populations was confirmed by GWAS. The most significant SNPs associated with race 1 resistance were located within the region of *qAnt*1.*Lc*-3. The GWAS SNPs fine mapped the *qAnt*1.*Lc*-3 region to 1.6 Mb containing candidate resistance genes.

A total of 57 candidate genes involved in plant defense against biotic and abiotic stress are predicted within the region of *qAnt*1.*Lc*-3 identified in the biparental mapping (Supplemental Table S4). Among these, 13 are NB-LRR class R genes, and 14 are transmembrane protein (TM) genes known as ‘oth-R’ genes (Sanseverino and Ercolano, 2012; Sekhwal et al., 2015). In plant genomes, the NB-LRR class R genes are abundant and often clustered on specific chromosomes due to tandem and segmental duplications (Leister, 2004). Congruently, most of the annotated NBS-LRR genes in the lentil genome are located on chromosome 3 (Ramsay et al., *unpublished data*). Thus, a cluster of candidate R genes identified within the region of *qAnt*1.*Lc*-3 could possibly account for improved resistance to race 1 in lentil. Transmembrane (TM) proteins are part of a plant cell complex membrane-associated receptors that mediate signal transduction between the extra- and intracellular environments in defense system and are mainly involved in conferring a broader resistance spectrum in plants, including the known *Mlo* gene (Büschges et al., 1997; Xiao et al., 2001; Brandwagt et al., 2002; Ma et al.,2014). The most significant SNP markers from GWAS analysis also identified genes involved with cellulose synthase and anthranilate N-benzoyltransferase protein within *qAnt*1.*Lc*-3. Cellulose synthases play an important role in mediating cell wall changes in the epidermal layers in response to defense against pathogens (Douchkov et al., 2016). Anthranilate N-benzoyltransferase genes are important in phytoalexins biosynthesis, which provide enhanced protections against pathogens (Grayer and Kokubun, 2001).

In conclusion, we combined the use of biparental QTL mapping and GWAS to identify QTL and candidate genes for anthracnose race 1 resistance in lentil. The major effect QTL (qAnt1.Lc-3) identified on chromosome 3, explained 66.6 to 69.8% of the phenotypic variance and was confirmed via GWAS analysis. Across the genome, GWAS identified 14 SNPs associated with race 1 resistance. The SNP markers identified that were associated with the candidate genes can be used for MAS to advance molecular breeding approaches for improving anthracnose resistance in lentil.

## SUPPLEMENTAL MATERIAL

Four supplemental tables (Supplemental Table S1 – S4) and three supplemental figures (Supplemental Figure S1-S3) are included as supporting information.

## ACKNOWLEDGEMENTS

The authors gratefully acknowledge funding from the Natural Sciences and Engineering Research Council of Canada (NSERC) Industrial Research Chair Program, the Saskatchewan Pulse Growers, and the University of Saskatchewan. We are grateful for financial support from a project ‘Application of Genomics to Innovation in the Lentil Economy (AGILE)’ funded by Genome Canada, Saskatchewan Pulse Growers, Western Grains Research Foundation, and the Province of Saskatchewan, and managed by Genome Prairie. We are also thankful for the technical assistance of the bioinformatics, molecular and pathology lab staff of the Pulse Breeding and Genetics group at the University of Saskatchewan.

## CONFLICTS OF INTEREST

The authors declare that they have no competing interests.

## AUTHOR CONTRIBUTIONS

TSG generated phenotypic data, performed data analyses, and wrote the manuscript. LR conducted SNP calling, TAH constructed the genetic map (LR-01), AV and KEB participated in experimental design and reviewed the manuscript. All authors read and approved the submitted version.

## DATA AVAILABILITY

The raw genotypic data supporting this study are available at https://knowpulse.usask.ca/Geneticmap/2686922; https://knowpulse.usask.ca/Geneticmap/2695342 and https://knowpulse.usask.ca/Lentil-AGILE-project or from the authors upon request. The SNPs that showed significant associations with anthracnose race 1 resistance are provided in Supplemental Information.

## Notes

### Competing Interest Statement

The authors have declared no competing interest.

