## Supplementary Material for "Identification of anthracnose (*Colletotrichum lentis*) race 1 resistance loci in lentil by integrating linkage mapping and a genome-wide association study"

**Supplemental Table S1.** Segregation of anthracnose race 1 in LR-01 and LR-18 populations, $\chi$ 2 test for 1:1 Mendelian ratio and corresponding probability.

| Population | Resistant | Susceptible | Total[#] | $\chi^2$ 1:1 | P |
|---|---|---|---|---|---|
| LR-01 | 47 | 55 | 102 | 0.63 | 0.428 |
| LR-18 | 58 | 80 | 138 | 3.51 | 0.061 |

[#]one RIL line showed a heterozygous reaction was not included in LR-18 population

**Supplemental Table S2.** Summary statistics of the lentil LR-01 (ILL 1704 × CDC Robin) population genetic linkage map.

| Linkage groups | Number of SNP markers | Number of independent marker loci[¥] | Numbers of BIN markers | Number of singleton markers | Map length (cM) | Average marker interval (cM) | Maximu m gap (cM) |
|---|---|---|---|---|---|---|---|
| LG1 | 2806 | 337 | 104 | 233 | 200.6 | 0.6 | 4.7 |
| LG2 | 5120 | 392 | 124 | 268 | 250.1 | 0.6 | 3.5 |
| LG3 | 4540 | 442 | 146 | 296 | 299.5 | 0.7 | 6.5 |
| LG4 | 3563 | 484 | 158 | 326 | 271.2 | 0.6 | 4.8 |
| LG5 | 2004 | 333 | 115 | 218 | 197.4 | 0.6 | 3.3 |
| LG6 | 1891 | 445 | 169 | 276 | 260.5 | 0.6 | 2.9 |
| LG7 | 1710 | 295 | 105 | 190 | 164.6 | 0.6 | 2.9 |
| Total | 21634 | 2728 | 921 | 1807 | 1643.9 | | |

[¥] Number of independent marker loci includes the number of BIN markers and number of singletons

**Supplemental Table S3.** SNP markers significantly associated with anthracnose race 1 resistance identified from trials in the growth chamber and polyhouse, and a combined lsmean of disease severity in a set of 200 lentil accessions.

| Environment | SNP Marker | Chr | Position (Mb)[#] | *P*.value | MAF | $R^{2\$}$ |
|---|---|---|---|---|---|---|
| **Combined** | Lcu.2RBY.Chr3.33827173 | 3 | 33827173 | 1.38E-06 | 0.16 | 0.65 |
| | Lcu.2RBY.Chr3.33827185 | 3 | 33827185 | 2.43E-08 | 0.15 | 0.67 |
| | Lcu.2RBY.Chr3.34117023 | 3 | 34117023 | 2.47E-10 | 0.14 | 0.69 |
| | Lcu.2RBY.Chr3.35384298 | 3 | 35384298 | 3.24E-06 | 0.14 | 0.65 |
| | Lcu.2RBY.Chr3.341261994 | 3 | 3.41E+08 | 4.30E-07 | 0.23 | 0.66 |
| | Lcu.2RBY.Chr3.417940994 | 3 | 4.18E+08 | 6.13E-06 | 0.06 | 0.64 |
| | Lcu.2RBY.Chr4.442702129 | 4 | 4.43E+08 | 8.32E-08 | 0.11 | 0.66 |
| | Lcu.2RBY.Chr4.442702133 | 4 | 4.43E+08 | 8.48E-08 | 0.11 | 0.66 |
| | Lcu.2RBY.Chr5.28582530 | 5 | 28582530 | 3.74E-06 | 0.12 | 0.65 |
| | Lcu.2RBY.Chr5.28637458 | 5 | 28637458 | 3.74E-06 | 0.12 | 0.65 |
| | Lcu.2RBY.Chr5.33721990 | 5 | 33721990 | 1.82E-09 | 0.21 | 0.68 |
| | Lcu.2RBY.Chr5.437910070 | 5 | 4.38E+08 | 3.23E-06 | 0.13 | 0.65 |
| | Lcu.2RBY.Chr5.437944230 | 5 | 4.38E+08 | 3.95E-06 | 0.13 | 0.65 |
| | Lcu.2RBY.Chr6.374326758 | 6 | 3.74E+08 | 8.88E-07 | 0.11 | 0.65 |
| **Growth chamber** | Lcu.2RBY.Chr2.36606480 | 2 | 36606480 | 4.64E-06 | 0.13 | 0.61 |
| | Lcu.2RBY.Chr2.57570472 | 2 | 57570472 | 3.08E-06 | 0.07 | 0.62 |
| | Lcu.2RBY.Chr2.59440536 | 2 | 59440536 | 3.08E-06 | 0.07 | 0.62 |
| | Lcu.2RBY.Chr2.72471727 | 2 | 72471727 | 4.81E-06 | 0.07 | 0.61 |
| | Lcu.2RBY.Chr3.29754683 | 3 | 29754683 | 2.86E-06 | 0.22 | 0.62 |
| | Lcu.2RBY.Chr3.33827173 | 3 | 33827173 | 5.27E-07 | 0.16 | 0.63 |
| | Lcu.2RBY.Chr3.33827185 | 3 | 33827185 | 1.05E-08 | 0.15 | 0.65 |
| | Lcu.2RBY.Chr3.34117023 | 3 | 34117023 | 1.86E-10 | 0.14 | 0.67 |
| | Lcu.2RBY.Chr3.35384298 | 3 | 35384298 | 4.15E-06 | 0.14 | 0.62 |
| | Lcu.2RBY.Chr3.57041703 | 3 | 57041703 | 5.91E-06 | 0.06 | 0.61 |
| | Lcu.2RBY.Chr3.162869024 | 3 | 1.63E+08 | 4.47E-06 | 0.17 | 0.62 |
| | Lcu.2RBY.Chr3.341261994 | 3 | 3.41E+08 | 1.46E-07 | 0.23 | 0.63 |

| | | | | | | |
|---|---|---|---|---|---|---|
| | Lcu.2RBY.Chr3.348840074 | 3 | 3.49E+08 | 2.34E-06 | 0.07 | 0.62 |
| | Lcu.2RBY.Chr3.349773406 | 3 | 3.5E+08 | 4.77E-07 | 0.06 | 0.63 |
| | Lcu.2RBY.Chr3.355843750 | 3 | 3.56E+08 | 5.28E-06 | 0.08 | 0.61 |
| | Lcu.2RBY.Chr3.400615790 | 3 | 4.01E+08 | 2.57E-06 | 0.06 | 0.62 |
| | Lcu.2RBY.Chr3.405224715 | 3 | 4.05E+08 | 2.98E-06 | 0.08 | 0.62 |
| | Lcu.2RBY.Chr3.417940994 | 3 | 4.18E+08 | 2.03E-06 | 0.06 | 0.62 |
| | Lcu.2RBY.Chr4.442702129 | 4 | 4.43E+08 | 3.27E-07 | 0.11 | 0.63 |
| | Lcu.2RBY.Chr4.442702133 | 4 | 4.43E+08 | 4.95E-07 | 0.11 | 0.63 |
| | Lcu.2RBY.Chr5.28377874 | 5 | 28377874 | 5.71E-06 | 0.31 | 0.61 |
| | Lcu.2RBY.Chr5.33721990 | 5 | 33721990 | 2.56E-09 | 0.21 | 0.65 |
| | Lcu.2RBY.Chr5.427535882 | 5 | 4.28E+08 | 4.50E-06 | 0.21 | 0.62 |
| | Lcu.2RBY.Chr5.430533888 | 5 | 4.31E+08 | 5.32E-06 | 0.12 | 0.61 |
| | Lcu.2RBY.Chr5.437910070 | 5 | 4.38E+08 | 3.14E-06 | 0.13 | 0.62 |
| | Lcu.2RBY.Chr5.437944230 | 5 | 4.38E+08 | 3.90E-06 | 0.13 | 0.62 |
| **Polyhouse** | Lcu.2RBY.Chr2.88737442 | 2 | 88737442 | 2.41E-06 | 0.07 | 0.59 |
| | Lcu.2RBY.Chr2.96273445 | 2 | 96273445 | 2.66E-06 | 0.12 | 0.59 |
| | Lcu.2RBY.Chr3.33827185 | 3 | 33827185 | 6.83E-07 | 0.15 | 0.59 |
| | Lcu.2RBY.Chr3.34117023 | 3 | 34117023 | 3.93E-08 | 0.14 | 0.61 |
| | Lcu.2RBY.Chr3.35384298 | 3 | 35384298 | 5.39E-06 | 0.14 | 0.58 |
| | Lcu.2RBY.Chr3.174824501 | 3 | 1.75E+08 | 4.58E-06 | 0.12 | 0.58 |
| | Lcu.2RBY.Chr4.13724159 | 4 | 13724159 | 3.55E-06 | 0.08 | 0.58 |
| | Lcu.2RBY.Chr4.442702129 | 4 | 4.43E+08 | 8.53E-07 | 0.11 | 0.59 |
| | Lcu.2RBY.Chr4.442702133 | 4 | 4.43E+08 | 9.07E-07 | 0.11 | 0.59 |
| | Lcu.2RBY.Chr5.33721990 | 5 | 33721990 | 1.49E-07 | 0.21 | 0.60 |
| | Lcu.2RBY.Chr6.374326758 | 6 | 3.74E+08 | 6.89E-07 | 0.11 | 0.59 |

#physical position, $explained phenotypic variance per marker

**Supplemental Table S4.** Potential **c**andidate resistance genes associated with anthracnose race 1 resistance in the interval of the QTL detected in two biparental RIL populations: LR-01 (ILL 1704 × CDC Robin) and LR-18 (CDC Robin × 964a-46), and GWAS regions according to gene annotation of lentil reference genome (v2.0; https://knowpulse.usask.ca/genome-assembly/Lc.2RBY)

| Chr[#] | Start (bp) | End (bp) | Gene ID | Annotation |
|---|---|---|---|---|
| Chr3 | 30099204 | 30101106 | Lcu.2RBY.3g005210 | Thylakoid lumenal 19 kDa protein |
| Chr3 | 30175439 | 30179402 | Lcu.2RBY.3g005220 | Makorin RING finger protein |
| Chr3 | 30217795 | 30227923 | Lcu.2RBY.3g005240 | Makorin RING-zinc-finger protein |
| Chr3 | 30300254 | 30301749 | Lcu.2RBY.3g005250 | Anthocyanin 5-aromatic acyltransferase |
| Chr3 | 30457815 | 30458393 | Lcu.2RBY.3g005280 | Transmembrane protein |
| Chr3 | 30481674 | 30482215 | Lcu.2RBY.3g005290 | Transmembrane protein |
| Chr3 | 30491797 | 30492000 | Lcu.2RBY.3g005300 | Transmembrane protein |
| Chr3 | 30641401 | 30646173 | Lcu.2RBY.3g005310 | NB-ARC domain disease resistance protein |
| Chr3 | 30704762 | 30705187 | Lcu.2RBY.3g005330 | Transmembrane protein |
| Chr3 | 30735452 | 30735939 | Lcu.2RBY.3g005340 | Tubby C 2 protein |
| Chr3 | 30735946 | 30736590 | Lcu.2RBY.3g005350 | Wall-associated receptor kinase protein |
| Chr3 | 30738028 | 30738585 | Lcu.2RBY.3g005360 | Transmembrane protein |
| Chr3 | 30832778 | 30837304 | Lcu.2RBY.3g005390 | Transmembrane protein |
| Chr3 | 31031941 | 31039406 | Lcu.2RBY.3g005400 | Transmembrane protein |
| Chr3 | 31113487 | 31114027 | Lcu.2RBY.3g005460 | Transmembrane protein |
| Chr3 | 31118273 | 31118672 | Lcu.2RBY.3g005480 | Transmembrane protein |
| Chr3 | 31195851 | 31196232 | Lcu.2RBY.3g005530 | Transmembrane protein |
| Chr3 | 31698296 | 31700380 | Lcu.2RBY.3g005650 | Anthranilate phosphoribosyltransferase protein |
| Chr3 | 31731103 | 31731724 | Lcu.2RBY.3g005660 | ORF1 |
| Chr3 | 32321603 | 32324284 | Lcu.2RBY.3g005730 | Myb transcription factor |
| Chr3 | 32693429 | 32694293 | Lcu.2RBY.3g005760 | Ulp1 protease family, carboxy-terminal domain |
| Chr3 | 32824221 | 32824963 | Lcu.2RBY.3g005770 | Ulp1 protease family, carboxy-terminal domain |
| Chr3 | 32892575 | 32892931 | Lcu.2RBY.3g005790 | Subtilisin-like serine protease |
| Chr3 | 32993043 | 32999825 | Lcu.2RBY.3g005810 | Multidrug and toxic compound extrusion protein |
| Chr3 | 33110296 | 33115291 | Lcu.2RBY.3g005840 | ARM repeat CCCH-type zinc finger protein |
| Chr3 | 33694640 | 33695284 | Lcu.2RBY.3g005860 | Anthranilate N-benzoyltransferase |
| Chr3 | 33819256 | 33828131 | Lcu.2RBY.3g005880 | Cellulose synthase |
| Chr3 | 33840742 | 33843043 | Lcu.2RBY.3g005900 | F-box SKIP23-like protein |

| | | | | |
|---|---|---|---|---|
| Chr3 | 34117126 | 34118497 | Lcu.2RBY.3g005910 | Anthranilate N-benzoyltransferase |
| Chr3 | 34413587 | 34439978 | Lcu.2RBY.3g005930 | Ulp1 protease family, carboxy-terminal domain |
| Chr3 | 34608235 | 34609591 | Lcu.2RBY.3g005950 | Anthranilate N-benzoyltransferase |
| Chr3 | 34618281 | 34620298 | Lcu.2RBY.3g005960 | Myb/SANT-like DNA-binding domain protein |
| Chr3 | 34650454 | 34651468 | Lcu.2RBY.3g005970 | Ulp1 protease family, carboxy-terminal domain |
| Chr3 | 34722504 | 34723667 | Lcu.2RBY.3g005990 | F-box SKIP23-like protein |
| Chr3 | 34936710 | 34937891 | Lcu.2RBY.3g006020 | F-box SKIP23-like protein |
| Chr3 | 35204566 | 35208726 | Lcu.2RBY.3g006030 | Polygalacturonase |
| Chr3 | 35383081 | 35387584 | Lcu.2RBY.3g006090 | Disease resistance protein /TIR-NBS-LRR class) |
| Chr3 | 35415534 | 35417750 | Lcu.2RBY.3g006110 | Zinc finger/RING finger family protein |
| Chr3 | 35783673 | 35784401 | Lcu.2RBY.3g006240 | MADS-box transcription factor |
| Chr3 | 35786719 | 35787457 | Lcu.2RBY.3g006260 | Zinc finger, C3HC4 type /RING finger protein |
| Chr3 | 35891715 | 35892401 | Lcu.2RBY.3g006280 | Zinc finger, C3HC4 type /RING finger protein |
| Chr3 | 35897123 | 35904908 | Lcu.2RBY.3g006300 | DNA topoisomerase II |
| Chr3 | 35972988 | 35973704 | Lcu.2RBY.3g006330 | LRR & NB-ARC domain disease resistance protein |
| Chr3 | 35974000 | 35976197 | Lcu.2RBY.3g006340 | LRR & NB-ARC domain disease resistance protein |
| Chr3 | 35976249 | 35981698 | Lcu.2RBY.3g006350 | LRR & NB-ARC domain disease resistance protein |
| Chr3 | 35981709 | 35982218 | Lcu.2RBY.3g006360 | NB-ARC domain disease resistance protein |
| Chr3 | 35982494 | 35982865 | Lcu.2RBY.3g006370 | CC-NBS-LRR resistance protein, putative |
| Chr3 | 36028314 | 36032907 | Lcu.2RBY.3g006380 | LRR & NB-ARC domain disease resistance protein |
| Chr3 | 36037105 | 36044777 | Lcu.2RBY.3g006390 | LRR & NB-ARC domain disease resistance protein |
| Chr3 | 36891988 | 36892982 | Lcu.2RBY.3g006490 | Transmembrane protein |
| Chr3 | 36920159 | 36920804 | Lcu.2RBY.3g006500 | Transmembrane protein |
| Chr3 | 37677606 | 37684302 | Lcu.2RBY.3g006580 | PPR containing plant-like protein |
| Chr3 | 38288475 | 38295878 | Lcu.2RBY.3g006660 | LRR & NB-ARC domain disease resistance protein |
| Chr3 | 38538802 | 38539297 | Lcu.2RBY.3g006720 | Transmembrane protein |
| Chr3 | 38755826 | 38764156 | Lcu.2RBY.3g006750 | LRR & NB-ARC domain disease resistance protein |
| unitig0289 | 1917198 | 1921007 | Lcu.2RBY.L001220 | LRR & NB-ARC domain disease resistance protein |
| unitig0289 | 1939874 | 1942161 | Lcu.2RBY.L001240 | NB-ARC domain disease resistance protein |

[#] Chromosome

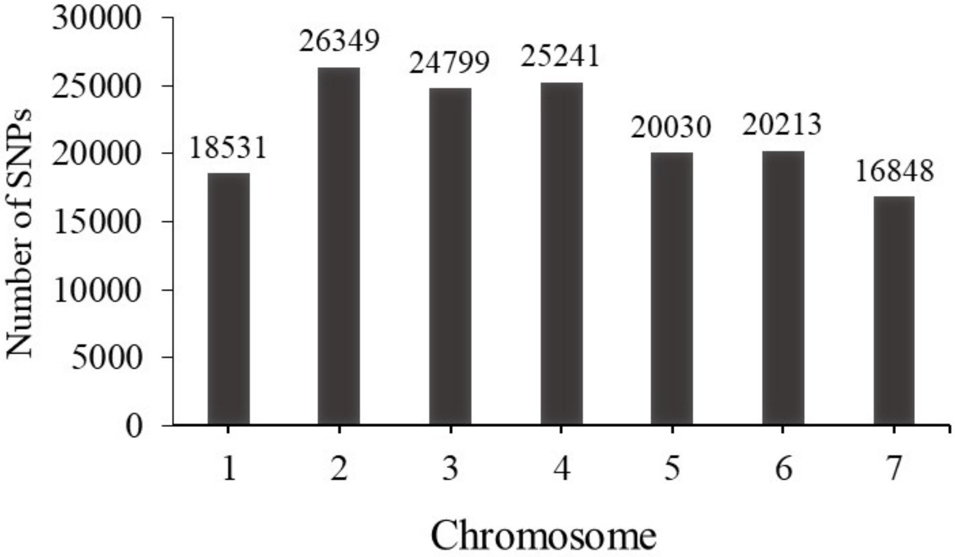


**Supplemental Figure S1.** Summary of SNP markers per chromosome used for GWAS analysis

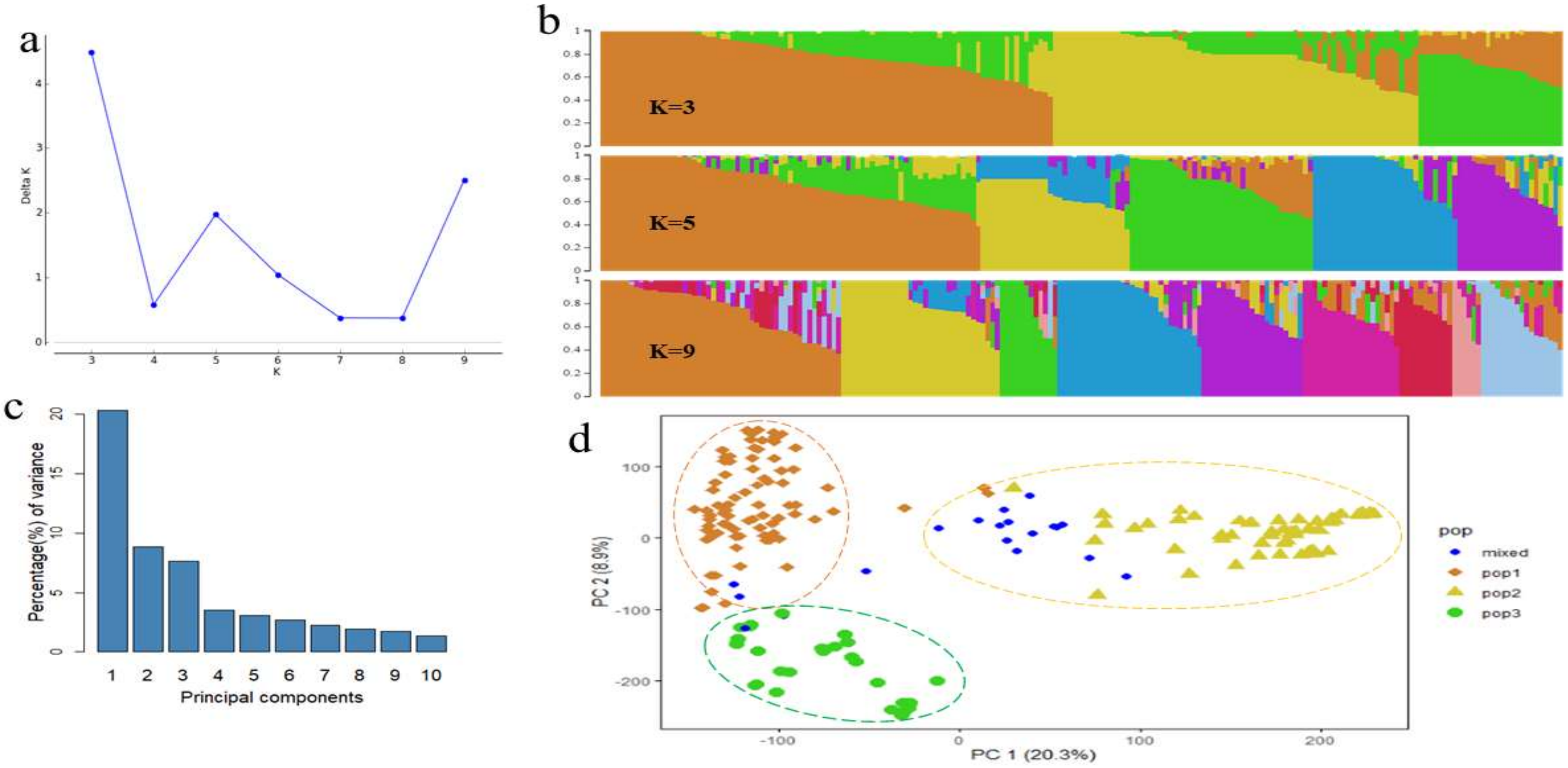


**Supplemental Figure S2.** The population structure of 200 lentil accessions was identified by the STRUCTURE admixture model and principal component analysis (PCA), which were then used for GWAS analysis. (**a**) delta K values, (**b**) population structure for models with K = 3, K=5 and K = 9, each genotype is represented by a vertical line, (**c**) percent of the variation explained by the first ten principal components, (**d**) scattered plot of the first and second principal components. The PCA plot is colored based on subpopulations (K=3) from the admixture model, whereas the blue dots represent genotypes with estimated membership fraction <60% and assigned as a mixed population.

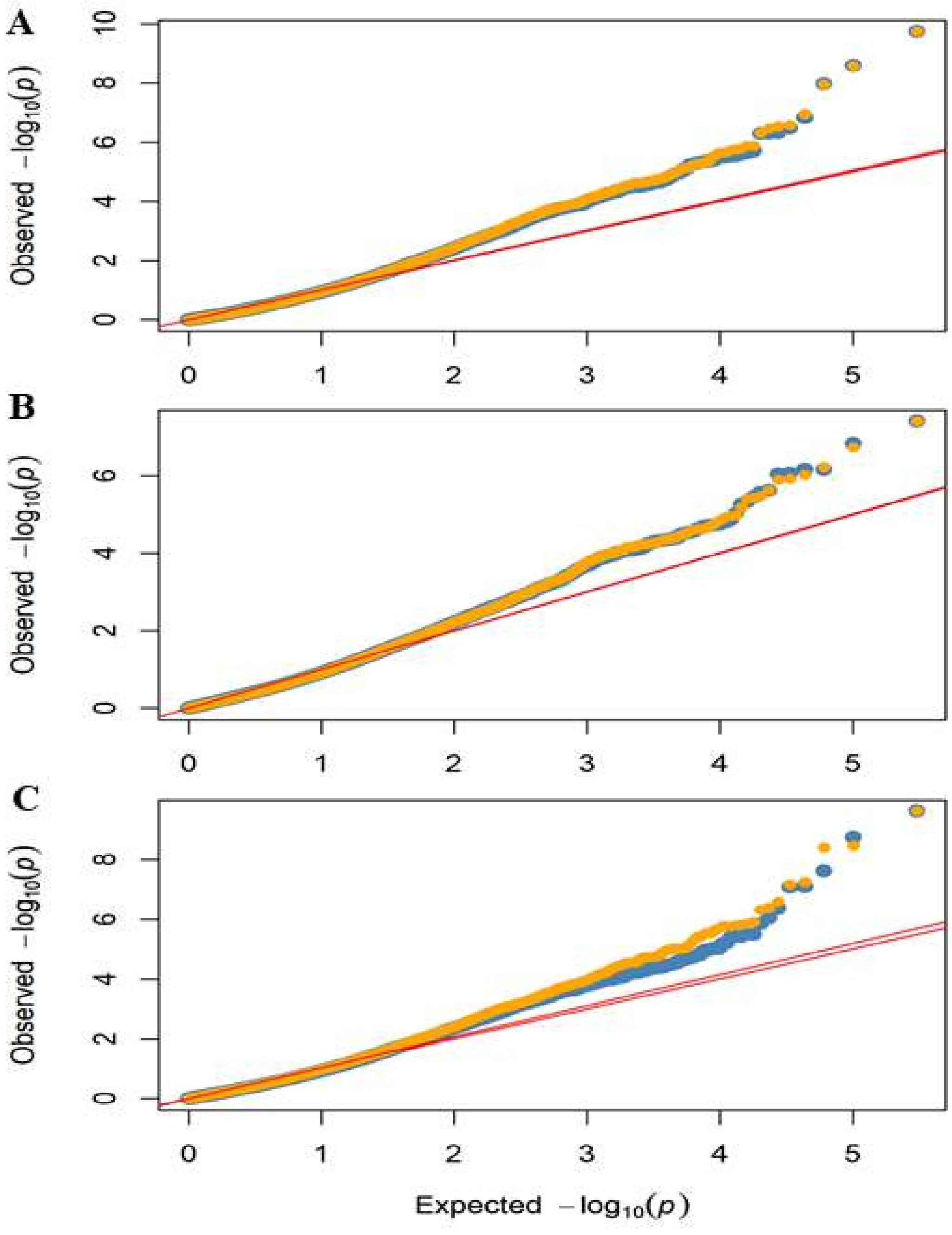


**Supplemental Figure S3.** Quantile-quantile (Q-Q) plots comparing the distribution of observed versus expected p-values for genome-wide association study of 200 lentil accessions evaluated for anthracnose race 1 severity using mixed linear model (MLM) analysis in the: A) growth chamber and B) polyhouse, and C) the combined lsmean from both environments. Orange dots represent the MLM approach using population structure (K=3) and kinship matrices, and the blue dots represent the model for principal component (PC=3) and kinship
